# GPMAW Glyco-Search: An Integrated Workflow for Identification and Validation of Intact Sialylated N-Glycopeptides

**DOI:** 10.64898/2026.08.05.743002

**Authors:** Maria K. Petersen, Simon N. Mule, Sara E. Lendal, Arkadiusz Nawrocki, Giuseppe Palmisano, Peter Højrup, Martin R. Larsen

## Abstract

Comprehensive analysis of intact sialylated N-glycopeptides remains challenging because of their low abundance, extensive structural heterogeneity, and limited peptide backbone fragmentation during tandem mass spectrometry. Here, we present an integrated workflow for high-confidence identification of intact sialylated N-glycopeptides that combines selective TiO₂ enrichment, dual LC-MS/MS analysis of intact and deglycosylated glycopeptides, and the GPMAW glyco-search platform based on high-accuracy mass mapping. Unlike most conventional glycoproteomics search engines, GPMAW uses experimentally identified deglycopeptides to constrain glycan assignment before matching intact glycopeptide precursor masses to candidate glycan compositions. Identifications were validated using diagnostic oxonium ions, glycopeptide-associated Y-ion fragments, and an experimentally derived glycopeptide score. In addition, GPMAW integrates an interactive spectrum annotation interface that enables rapid manual validation of candidate identifications through visualization of annotated Y-ion series, oxonium ions, and peptide fragments, allowing individual assignments to be readily accepted or rejected.

The workflow was optimized using bovine fetuin, validated on standard glycoproteins, and applied to depleted human plasma, where more than 2800 unique intact sialylated N-glycopeptides were identified across hundreds of glycosites and glycoproteins. Moreover, more than 1000 unique N-glycopeptides were identified from only 1 μL of plasma. Comparative analysis demonstrated that GPMAW glyco-search identified more confidently assigned intact sialylated N-glycopeptides than three widely used N-glycoproteomics search engines while maintaining high reproducibility and low false-positive rates following manual validation. Together, this workflow provides a robust, flexible, and accessible platform for large-scale, high-confidence characterization of intact N-glycopeptides and establishes experimentally constrained glycan composition assignment combined with interactive spectrum validation as an effective strategy for reducing ambiguity in N-glycoproteomics.

**Highlights:**

- The program “GPMAW glyco-search” enables high-accuracy mass mapping for confident identification of intact N-glycopeptides.
- Integrated workflow combining TiO₂ enrichment, dual LC-MS/MS of intact and deglycosylated glycopeptides and GPMAW glyco-search for intact sialylated N-glycopeptides.
- Optimized TiO₂ enrichment provides >95% selective enrichment of sialylated N-glycopeptides from complex biological samples.
- Interactive spectrum annotation and Y-ion-based scoring enable rapid manual validation and high-confidence glycopeptide identification.
- GPMAW glyco-search confidently identified more intact sialylated N-linked glycopeptides compared to three established glycoproteomics search engines.

## Introduction

Protein glycosylation is a fundamental co-/post-translational modification essential for numerous biological processes [1, 2]. N-glycosylation, the co-translational covalent attachment of oligosaccharides to asparagine (N) residues, typically occurs at the consensus motif N-X-S/T/C (X ≠ proline) [3–7]. Because glycan biosynthesis is not template-driven, individual glycosylation sites (glycosites) often exhibit extensive heterogeneity, posing major challenges for comprehensive glycoprotein characterization [8].

Glycoproteomics and glycomics are complementary approaches to globally profile protein glycosylation. Glycomics entails comprehensive structural analysis of glycans released from glycoproteins [2, 9, 10]. This approach defines the repertoire of glycans in a biological system but loses information about the protein carriers and specific glycosites. Glycoproteomics, in contrast, analyses intact glycopeptides to preserve linkage between glycans and peptides.

Early glycoproteomic workflows often removed N-glycans enzymatically (e.g., with peptide N-glycosidase F (PNGase F)) prior to mass spectrometry (MS) identification [11–13]. This enabled identification of formerly N-glycosylated peptides (marked by an N-to-D conversion after deglycosylation) [14] but discarded the glycans, thereby losing glycoform-specific information and could complicate biological interpretation. Recent advances have focused on liquid chromatography-tandem MS (LC-MS/MS) analysis of intact glycopeptides, which retains the glycan attached to the peptide thereby revealing the glycoprotein identity, glycosite, and glycan composition in a single experiment. However, intact glycopeptides are often low in abundance, prone to ionization suppression [15], and fragment inefficiently in MS/MS (especially when sialylated) [16–18], making their intact analysis difficult and challenging.

Selective enrichment of glycopeptides is commonly employed to improve identification. Strategies include hydrazide chemistry [19], hydrophilic interaction liquid chromatography (HILIC) [20–22], lectin affinity capture [23], and titanium dioxide (TiO_2_) chromatography [24]. The latter is particularly effective for selective enrichment of sialylated glycopeptides [24, 25]. For MS/MS fragmentation, collision-induced dissociation (CID) and higher-energy collisional dissociation (HCD) that preferentially cleave glycosidic bonds, yielding abundant glycan fragment ions but limited peptide fragments [26, 27], are used. In contrast, electron-capture dissociation (ECD) [28] and electron-transfer dissociation (ETD) [29] cleave peptide backbones while largely preserving glycans, although both perform poorly on glycopeptides carrying sialic acids due to inefficient backbone fragmentation (CID/HCD) or extensive charge reduction (ETD) [16].

Numerous bioinformatic tools have been developed for intact N-glycopeptide identification, and a recent HUPO community study benchmarked many of these search engines on serum glycopeptide data [30]. Existing algorithms generally employ glycan-first, peptide-first, or hybrid search strategies. Glycan-first approaches identify glycans from diagnostic fragment ions before assigning the peptide (e.g., pGlyco [31], Sweet-Heart [32]), whereas peptide-first approaches identify the peptide first and subsequently assign the glycan composition (e.g., Byonic [33], MSFragger [34]). Hybrid methods search peptide and glycan databases simultaneously (e.g., GlycoPAT [35], StrucGP [36]). Despite these advances, reliable identification of sialylated N-glycopeptides remains challenging because sialylated glycans often reduce peptide backbone fragmentation and increase glycan ambiguity [16]. To overcome these limitations, mass-difference approaches (e.g., GPQuest [37], ArMone 2.0 [38], and MAGIC [39]) infer glycan composition from the mass difference between an intact glycopeptide and its corresponding deglycosylated peptide, followed by validation using diagnostic glycan fragment ions [40–42].

Here, we present an integrated workflow for large-scale analysis of intact sialylated N-glycopeptides from simple and complex samples. Our strategy combines optimized TiO_2_-based enrichment of sialylated N-glycopeptides, high-accuracy LC-MS/MS, and a dedicated glyco-search module implemented in GPMAW (General Protein/Mass Analysis for Windows), a software platform for protein and peptide mass spectrometry [31]. Unlike many specialized glycoproteomics search engines, GPMAW runs on a standard Windows PC requiring just a standard license. The GPMAW glyco-search module identifies candidate glycan compositions by accurate mass mapping between intact glycopeptides and their corresponding PNGase F/A-deglycosylated peptides followed by validation using diagnostic glycan fragment ions and a Y-ion-based scoring system. The interactive interface further enables manual inspection of annotated spectra and flexible filtering of candidate identifications.

The workflow was optimized on the well-characterized N-glycoprotein fetuin and additional model glycoproteins before being applied to human plasma, where more than 2800 N-glycopeptides were identified from minute material. TiO_2_ chromatography enriched sialylated N-glycopeptides with > 94% specificity across all samples. Comparative analysis demonstrated that GPMAW glyco-search outperformed three established N-glycoproteomics search engines. Together, selective enrichment, optimized fragmentation, and dedicated informatics provide a robust platform for confident identification and validation of intact sialylated N-glycopeptides while substantially improving analytical depth.

## Material and Methods

Protein samples such as the standard proteins (all Sigma-Aldrich) bovine fetuin, human alpha-1-acid glycoprotein 1, human serotransferrin, human fibrinogen alpha/beta chain, bovine thyroglobulin, and human plasma were prepared similarly except for depletion and high pH (HpH), which were only performed on the depleted human plasma samples.

### Depletion of human plasma

A total of 40 µL human plasma was depleted of its 14 most abundant proteins using High Select^TM^ Top14 Abundant Protein Depletion Resin (Thermo Scientific) according to manufacturer’s protocol with minor modifications [32]. A total of 300 µL depletion resin was used per 10 µL plasma. The depletion was performed on MobiSpin Columns “F” (10µm filter, Boca Scientific Inc.) with 100mM ammonium bicarbonate (MP Biomedicals) as the buffer. The flowthrough containing the remaining proteins was lyophilized prior to proteolysis.

### Proteolytic digestion of standard proteins and plasma proteins

Protein samples were redissolved in 100mM HEPES (Sigma-Aldrich)/1% sodium deoxycholate (SDC, Sigma-Aldrich) buffer, pH 8.0. Proteins were reduced with 10 mM dithiothreitol (DTT, Sigma-Aldrich) for 30 minutes (min) at room temperature (RT), followed by alkylation with 20 mM iodoacetamide (IAA, Sigma Aldrich) for 30 min in the dark at RT. The reaction was quenched with 5 mM DTT for 10 min at RT. Proteins were digested overnight at 37°C with 3% w/w trypsin (methylated in-house [33]). After incubation, the trypsin was inactivated and SDC precipitated by adding 1% trifluoroacetic acid (TFA, Sigma-Aldrich) followed by centrifugation at 20.000 g for 10 min at RT. The supernatants were transferred to low binding Eppendorf tubes for further analysis.

### Enrichment of sialylated N-linked glycopeptides using TiO_2_ chromatography

Sialylated N-glycopeptides were enriched using batch-mode TiO_2_ chromatography, employing 1.0 mg of TiO_2_ beads (5µm, GL Sciences) per 100 µg of peptide sample according to previously published protocol [24]. Peptides were resuspended in 1mL of 1M glycolic acid (Sigma-Aldrich)/80% acetonitrile (ACN, VWR Chemicals)/5% TFA and incubated with the TiO_2_ beads for 10 min at RT with constant shaking. The supernatant was transferred to a fresh vial containing 0.5 mg TiO_2_ beads per 100 µg peptide sample for a second 10-min incubation with constant shaking. The TiO_2_ beads from both incubations were pooled and washed with 100 µL 80% ACN/1% TFA. After centrifugation and removal of the supernatant, the TiO_2_ beads were dried for 10 min. Intact sialylated N-glycopeptides were eluted with 0.1% triethylamine (TEA, Sigma-Aldrich), pH 11.4. The eluate was passed through a C8 filter-tip (3M Bioanalytical Technologies), which was subsequently washed with 10 µL 50% ACN in water and collected into the same sample. A small aliquot of each sample was analyzed by LC-MS/MS to confirm an efficiency of >90% enriched N-linked glycopeptides based on the m/z 204.09 and 366.14 diagnostic oxonium ions. The eluted N-glycopeptides were lyophilized prior to further analyses.

### High pH reversed phase (RP) fractionation

Purified sialylated N-glycopeptides from depleted plasma was dissolved in 30 µL solvent A (20 mM ammonium formate, pH 9.5, Sigma-Aldrich) and fractionated on an Acquity UPLC^®^- Class CSHTM C18 column (Waters) using a Dionex Ultimate 3000 HPLC system (Thermo Scientific). Separation was conducted with a 116-min gradient of solvent B (80% ACN, 20% solvent A) in solvent A (Solvent B: 2-50% in 60 min, 50-70% in 10 min, and 70-95% in 5 min) at a flow rate of 5 µL/min. Fractions were concatenated into 12 fractions with collection every 151 second and the samples were lyophilized prior to LC-MS/MS.

### Deglycosylation of sialylated N-glycopeptides

A fraction (30%) of sialylated N-glycopeptides purified by TiO_2_ from a tryptic digestion of the standard proteins, un-depleted human plasma and high pH RP fractions from depleted plasma was subsequently deglycosylated using PNGase A (New England Biolabs)(5000 U/mL), PNGase F (New England Biolabs)(500.000 U/mL) and sialidase A (Agilent Technologies)(5 U/mL) over-night in 100 mM HEPES, pH 7.5. The amount of deglycosylation enzymes used depended on the samples and amount of sialylated glycopeptides estimated in each sample. After deglycosylation, the samples were acidified, and the peptides were purified using an Oligo R3 RP micro-column as described previously [34]. The deglycosylated peptides were lyophilized prior to LC-MS/MS.

### Liquid Chromatography-Tandem Mass Spectrometry (LC-MS/MS)

Intact sialylated N-glycopeptides were dissolved in 0.05% TFA (Sigma-Aldrich) and separated by RP chromatography using either a nanoEASY-LC (Thermo Scientific) or a Vanquish NEO system (Thermo Scientific) coupled to an Orbitrap ECLIPSE Tribrid (Thermo Scientific). Intact sialylated N-glycopeptides from the standard proteins were loaded onto an in-house 2 cm pre-column (100 μm inner diameter packed with ReproSil-Pur C18 AQ 3 μm reversed-phase material (Dr. Maisch, Germany)) using the nanoEASY-LC system and intact sialylated

N-glycopeptides from plasma (depleted and non-depleted) were loaded onto a two-column system with a 5 mm x 300 µm Acclaim™ PepMap™ 100 C18 HPLC trap column (5 µm, Thermo Scientific) using the Neo LC system. For both LC systems the analytical column was an in-house packed 20 cm x 100 µm ID Reprosil-Pur C18-AQ RP analytic column (3 µm; Dr. Maisch GmbH, Germany).

The intact glycopeptides from the standard glycoproteins were separated using an increasing gradient of solvent B (95% ACN/0.1% formic acid (FA)) at a flow rate of 250-400 nL/min. The LC gradient for the standard glycoproteins using the nanoEASY-LC was 2-40%B in 30 min, 40-70%B in 1 min, 70%B for 3 min, 70-100%B in 1 min and 100%B for 3 min. The flow was 300 nL/min. Full MS1 scans were acquired with an automatic gain control (AGC) target of 250%, scan range at 700-2000 and an Orbitrap resolution of 240k. The most abundant peptide ions were selected in a 3 s cycle time from MS1 using activation by HCD fragmentation at stepped collision energy of 22% and 28% at 50k FWHM Orbitrap resolution. The AGC target was set to 500% and a maximum injection time of 100 ms. A dynamic exclusion of 20 seconds was used with an isolation window of 1.0 m/z.

Intact glycopeptides from the depleted plasma high pH RP fractions were separated using the Neo setup with the following LC gradient; 1-28%B in 45 min, 28-50%B in 15 min, 50-70%B in 1 min, 70%B for 3 min and 70-95%B in 1 min and 95%B for 5 min. The flow was 400 nL/min. Full MS1 scans were acquired with an automatic gain control (AGC) target of 250%, scan range at 800-2000 and an Orbitrap resolution of 240k. The most abundant peptide ions were selected in a 3 s cycle time from MS1 using activation by HCD fragmentation at stepped collision energy (sceHCD) of 25, 28 and 32% at 50k FWHM Orbitrap resolution. The AGC target was set to 500% and a maximum injection time of 150 ms. A dynamic exclusion of 20 seconds was used with an isolation window of 1.0 m/z.

The intact glycopeptides from non-depleted plasma were separated using the Neo setup with the following LC gradient; 1-28%B in 100 min, 28-50%B in 20 min, 50-70%B in 1 min, 70%B for 5 min and 70-95%B in 1 min and 95%B for 5 min. The settings for the MS/MS on the Eclipse were like the depleted plasma fractions.

The deglycopeptide fractions were dissolved in 5 µL 0.1% TFA and loaded onto a 2 cm precolumn (3 µm; Dr. Maisch GmbH, Germany) using a nanoEASY-LC (Thermo Scientific) coupled to an Orbitrap Astral (Thermo Scientific). The peptides were eluted onto a 20 cm analytical column (100 µm ID Reprosil-Pur C18-AQ RP analytic column (1.9 µm; Dr. Maisch GmbH, Germany)) using the following gradient: 2-40% in 20 min, 40-100% in 2 min, 100% for 5 min, and 100-2% in 1 min. Full MS1 scans were acquired in the Orbitrap with an automatic gain control (AGC) target of 500%, maximum injection time of 5 ms, scan range at 300-1200 and an Orbitrap resolution of 240k. The most abundant peptide ions were selected in a 1 s cycle time from MS1 for MS2 in the Astral analyzer using activation by HCD fragmentation at normalized collision energy of 30%, an AGC target of 500% and a maximum injection time of 10 ms. A dynamic exclusion of 7 seconds was used with an isolation window of 1 m/z.

The raw MS data, MGF files, and associated search files have been deposited to the ProteomeXchange Consortium via the PRIDE [35] partner repository with the dataset identifier PXD081982.

### Proteome Discoverer Analysis

Deglycosylated peptides were identified using Proteome Discoverer (PD, v2.5, Thermo Scientific) using the SEQUEST HT search engine, against the human UniProt protein database (v. 15.07.2025, 20574 entries). Peptides were searched with following fixed parameters: precursor mass tolerance 10 ppm, MS/MS mass tolerance 0.03 Da, trypsin as enzyme, maximum of 1 missed cleavage, and carbamidomethylation (C) as fixed modification. Variable modifications included deamidation (N). Only peptides with a q-value <0.01 (Percolator [36]), were considered for further analysis (≤1% false discovery rate). The identified deglycosylated peptides were filtered and validated in Excel and the list of formerly sialylated N-glycopeptides were saved in Excel 97-format (xls) for uploading into GPMAW (https://gpmaw.com/). Mgf files were generated from the intact sialylated N-glycopeptide raw data using the Spectrum Exporter node in PD.

### GPMAW glyco-search

Intact sialylated N-glycopeptides were assigned to their corresponding deglycosylated peptides using the GPMAW (v15.1) glyco-search tool through accurate mass mapping, followed by automated filtering and manual validation. For each deglycosylated peptide, GPMAW searched intact glycopeptide precursor masses from the MGF file within a user-defined precursor mass tolerance of 5–10 ppm and calculated all compatible glycan compositions. MS/MS information was imported directly from the corresponding MGF files and linked to each candidate glycopeptide using the scan number. Fragment ions were matched with an isotope mass tolerance of 0.05 Da. Candidate glycopeptides were subsequently filtered using characteristic fragment ions generated during HCD fragmentation of N-glycopeptides. Candidates lacking either the Peptide + HexNAc (Y1; peptide + 203.079 Da) fragment ion, the Peptide + Core (Y5; peptide + (HexNAc)2(Hex)3, peptide + 892.317 Da) fragment ion, or the HexNAc oxonium ion (m/z 204.087) were discarded. Duplicate assignments, based on identical combination of peptide sequences and glycan compositions, were removed by retaining the candidate with the highest Y1 fragment ion intensity. Candidates with a Y1 fragment ion peak intensity below 20k were excluded, as low-intensity Y1 ions were found to be associated with substantially reduced confidence during manual spectrum validation.

To further reduce false-positive identifications, GPMAW glyco-search calculates a confidence score based on the summed intensity of the assigned Y1–Y6 ions relative to the total fragment ion intensity within the mass range spanning Y1 to peptide + core + HexNAc (Y6) (described in **Supplementary File S1**). Scores range from 0 to 1000. Based on manual interpretation of several thousand intact N-glycopeptide MS/MS spectra, scores above 650 were considered very high-confidence assignments, whereas candidates scoring below 450 were consistently false positives. Candidate glycopeptides with scores between 450 and 650 were subjected to manual inspection and included several co-eluting intact N-glycopeptides.

Final validation was performed by manual *de novo* interpretation of glycan fragmentation using the GPMAW glyco-search annotation interface, which displays color-coded fragment ion annotations directly on the individual MS/MS spectra generated from the MGF files. The validated list of intact N-glycopeptides was exported to Excel for downstream analysis, including evaluation of glycan compositions, fucosylation patterns, and site-specific sialylated N-glycan heterogeneity.

A detailed guide on how to use and navigate the GPMAW glyco-search tool can be found in **Supplementary File S1.**

### Byonic search

Following LC-MS/MS analysis of the TiO_2_ enriched glycopeptides from 1 µL plasma, the resulting raw files were submitted to intact N-linked glycopeptide searches using the Byonic™ software (v2.10.47, Protein Metrics). For this analysis, the accession numbers from the 287 glycoproteins identified previously from the deglycoproteome analyses were loaded onto UniProt (https://www.uniprot.org/) and their corresponding protein sequences (FASTA format) downloaded for subsequent use as the protein database. The GPMAW glycan database, comprised of 282 unique glycans, was used here as the glycan database. Byonic™ searches were performed with the following modifications: precursor mass tolerance of 10 ppm, product ion mass tolerance of 0.05 Da, QTOF/HCD was set as the fragmentation method, carbamidomethylation of cysteine was set as the fixed modification, and a fully specific trypsin cleavage pattern allowing for 2 missed cleavages. Oxidation of methionine and glycosylation of asparagine were set as variable modifications. Precursor isotope by x was set to “Too high or low (narrow)”, which allows for the assigned precursors to be off the true precursor mass by -/+ 2 Da. Default maximum number of precursors per scan of 2 and smoothing width (m/z) 0.001 were used. A protein FDR was set at 0.01 (1%). The list of identified intact N-glycopeptides was filtered using a PEP2D cut-off < 0.001. Finally, non-redundant peptide-glycan sequence combinations were considered for subsequent comparisons.

### MSFragger search

For comparison, the same raw LC-MS/MS files were searched using the glyco-N-HCD workflow implemented in FragPipe computational platform v21.1 (https://fragpipe.nesvilab.org/). The MS/MS spectra were searched using the database search tool MSFragger v4.1 against the deglycoproteome protein sequence database previously described, which was appended with 50% of decoy sequences. A precursor-ion mass tolerance of 10 ppm and a fragment tolerance of 0.05 Da were set, with trypsin as the digestion enzyme limited to a maximum of 2 missed cleavages. Cysteine carbamidomethylation was specified as the fixed modification, while methionine oxidation (+15.9949) and N-terminal protein acetylation (+42.0106) were set as variable modifications. Similar to Byonic searches, the same glycan database from GPMAW was used. PeptideProphet was used for PSM validation, while PTM-Shepard was set to default settings for Glyco search (Glycan FDR < 0.01, glycan mass tolerance of 50 ppm). An isotope error range of −1 to +1 was used. The list of identified N-glycopeptides was filtered using a glycan q-value < 0.01, and the resulting high confident intact N-glycopeptides were filtered to remove redundancy prior to comparison analysis.

### Glyco-Decipher search

The same raw LC-MS/MS files were analyzed using Glyco-Decipher (v.1.0.2) [37] with the following settings: Similar to Byonic and MSFragger, a precursor and fragment mass tolerance of 10 ppm and 0.05 Da were used, respectively. Moreover, carbamidomethylation of cysteine and oxidation of methionine were used. Trypsin was set as the enzyme used to generate the peptides with full specificity, allowing for a maximum of 2 missed cleavages. The deglycoproteome protein database previously described was used, while the built-in GlyTouCan database [38], with 1755 unique N-glycan compositions, was used. SpectrumExpansion was activated to enable glycopeptide spectra matching of poorly fragmented peptide backbones [37]. A cut-off of glycan FDR <0.001 was applied to the identified intact glycopeptides from Glyco-Decipher searches, and the high confident intact N-glycopeptide identifications were filtered to remove peptide sequences and glycan composition redundancy for subsequent comparative analysis.

## Results and Discussion

### Integrated workflow for identification of intact sialylated N-glycopeptides

Our previous studies reported high efficiency for the selective enrichment of sialylated N-glycopeptides using TiO_2_ chromatography, when using optimized buffer conditions [24, 39]. Our workflow for identification of intact sialylated N-glycopeptides (**Figure 1**) relies on this selective enrichment using TiO_2_ chromatography combined with dual LC-MS/MS analysis of intact sialylated N-glycopeptides and their deglycosylated counterparts achieved through PNGase F/A and sialidase A treatment. The workflow can identify sialylated N-deglycosylated sites along with their potential sialylated glycan compositions in both simple and complex peptide samples. Notably, this workflow does not allow for the identification of individual linkages between monosaccharides (e.g., α2,3 or α2,6 sialylation).

**Figure 1:**
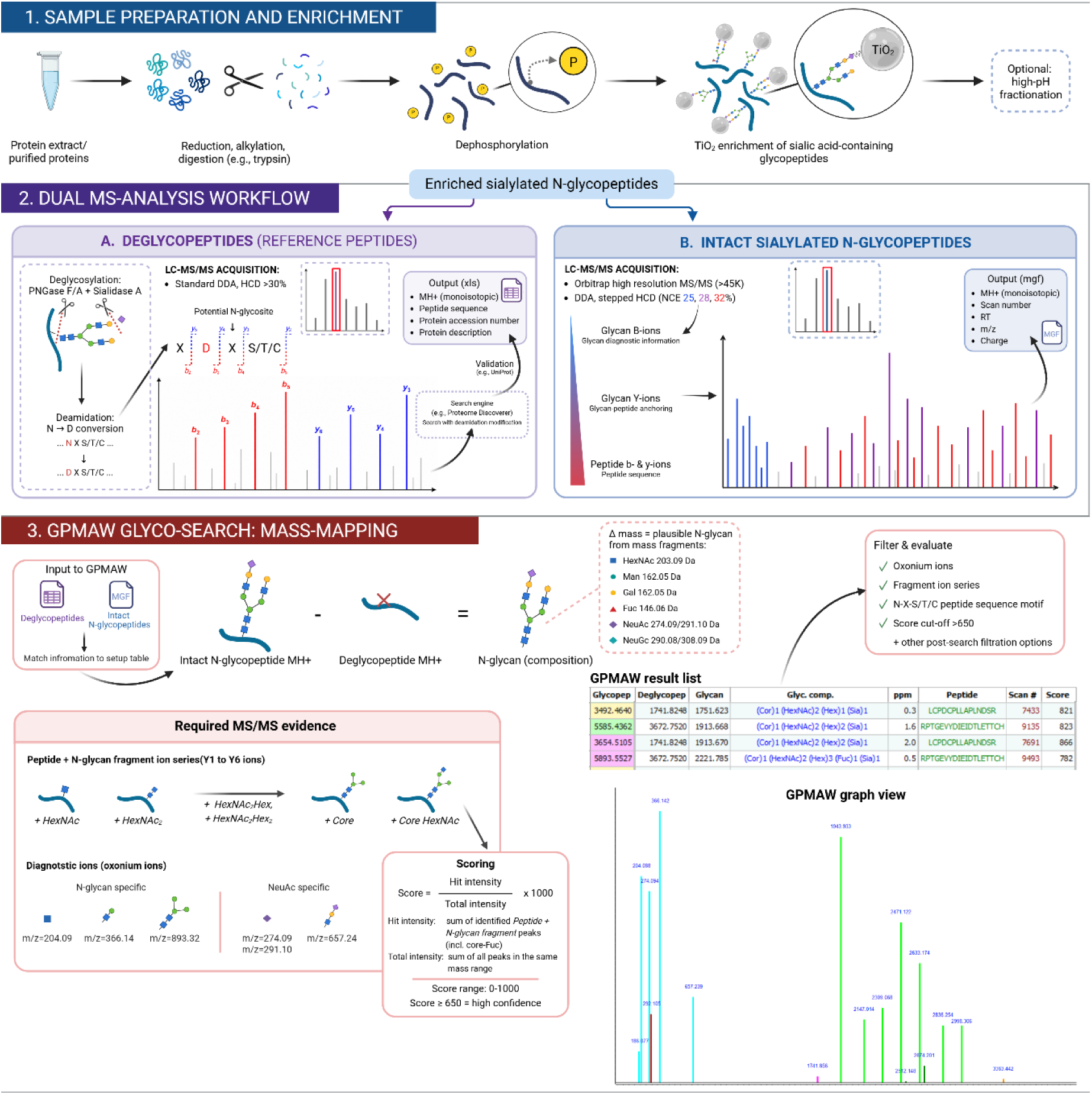
Integrated workflow for TiO₂ enrichment, dual LC-MS/MS analysis and GPMAW Glyco-search identification of intact sialylated N-glycopeptides. The workflow consists of three parts. **(1)** Sample preparation, digestion and TiO₂-based enrichment of sialylated N-glycopeptides, with optional high-pH reversed-phase fractionation of complex samples. **(2)** Dual LC-MS/MS analysis of enriched glycopeptides, including deglycosylation with PNGase F/A and sialidase A to identify N-glycosites and analysis of the remaining enriched sample as intact sialylated N-glycopeptides using stepped HCD fragmentation. **(3)** GPMAW Glyco-search integrates the intact glycopeptide and deglycopeptide datasets through accurate mass mapping of intact glycopeptides and corresponding deglycopeptides to assign candidate N-glycan compositions, followed by post-search validation based on diagnostic oxonium ions, glycan Y-ion series, peptide sequence motif, GPMAW score and manual inspection of annotated MS/MS spectra.

The LC-MS/MS raw data file generated after enzymatic deglycosylation of enriched sialylated N-glycopeptides is processed using PD, where formerly sialylated N-glycopeptides within the common consensus site N-X-S/T/C (N≠P) are identified by the deamidation “tag” originating from the hydrolysis from the enzymatic deglycosylation. The list of deglycopeptides is further validated and sorted for known N-glycosylation information from the UniProt database, as spontaneous deamidation can occur during sample preparation, especially in NG sites [40]. Next, an MGF file is generated in PD from the LC-MS/MS raw data file from the analysis of the intact sialylated N-glycopeptides using sceHCD fragmentation. This MGF file contains information on precursor and fragment ion m/z and intensities as well as retention time for the selected precursor ions. A glyco-search module implemented in GPMAW (https://www.gpmaw.com/) uses accurate mass mapping to assign glycan compositions to individual glycosites by aligning the masses of deglycosylated peptides with their corresponding intact sialylated N-glycopeptides from the MGF file. Candidate glycan compositions are calculated from the mass difference using the monosaccharide building blocks commonly found in mammalian N-glycans, including HexNAc, Hex, Fuc, and Neu5Ac. The non-human Neu5Gc can be included in the search but with restrictions (described in **Supplementary File S1**). As default, GPMAW glyco-search uses a glycan list consisting of 282 glycan compositions (**Supplementary Data S1 and described in Supplementary File S1**). However, any custom-made glycan list can be uploaded in the program. The built-in on-the-fly deconvolution algorithm should always be used for the best results, particularly as the calculation of the score is dependent on this. It can be turned off for testing purposes, or if the MGF file is already deconvoluted.

After high-accuracy mass mapping assignment of potential N-glycan compositions to individual deglycopeptides, a computational filtering is performed for characteristic signature fragment ions commonly observed in the fragmentation of N-glycopeptides, corresponding to the peptide plus the first attached GlcNAc (Y1), and the peptide plus an attached common N-glycan core structure (Y5), corresponding to the two GlcNAc linked to three mannoses. This filtering is based on previous observations that most N-glycopeptides, when fragmented with CID/HCD, yield intense Y1 [41] and Y5 ions. During filtering, the minimum intensity threshold for the Y1 fragment ion can be adjusted to increase the stringency of N-glycopeptide identification. This filtering step confirms the presence of N-glycopeptides and candidate matches lacking these signature fragment ions are discarded. Redundant assignments based on peptide sequence and assigned glycan composition are also filtered out. The GPMAW glyco-search interface enables detailed evaluation of fragment ion patterns through a built-in spectrum visualization tool, facilitating confirmation of N-glycopeptide assignments **(Figure 2A**). Furthermore, based on manual annotation of thousands of intact N-glycopeptide MS/MS spectra, we developed a score to allow for fast and accurate identification of N-linked glycopeptides in GPMAW glyco-search. The score is based on the empirical observation that, for correctly assigned N-glycopeptides, the majority of fragment ion intensity within the mass range spanning the Y1 to the Y6 ions originates from the correctly assigned peptide Y-ion series. The score is therefore calculated as the summed intensity of all assigned Y1–Y6 ions relative to the total fragment ion intensity within this mass range. An example and further explanation of the score is included in **Supplementary File S1**. Thus, any identified intact N-glycopeptide in GPMAW glyco-search is assigned a confidence score ranging from 1 to 1000. From the manual evaluation scores between 650 and 1000 are considered very high-confidence identifications whereas scores below 450 are considered false identifications. Co-isolation of two or more intact N-glycopeptides could result in slightly lower scores, and for obtaining slightly higher coverage of N-glycopeptides it is recommended that assigned glycopeptides with a score between 450 and 650 are manually inspected in the GPMAW glyco-search “Graph / evaluation” viewer.

**Figure 2:**
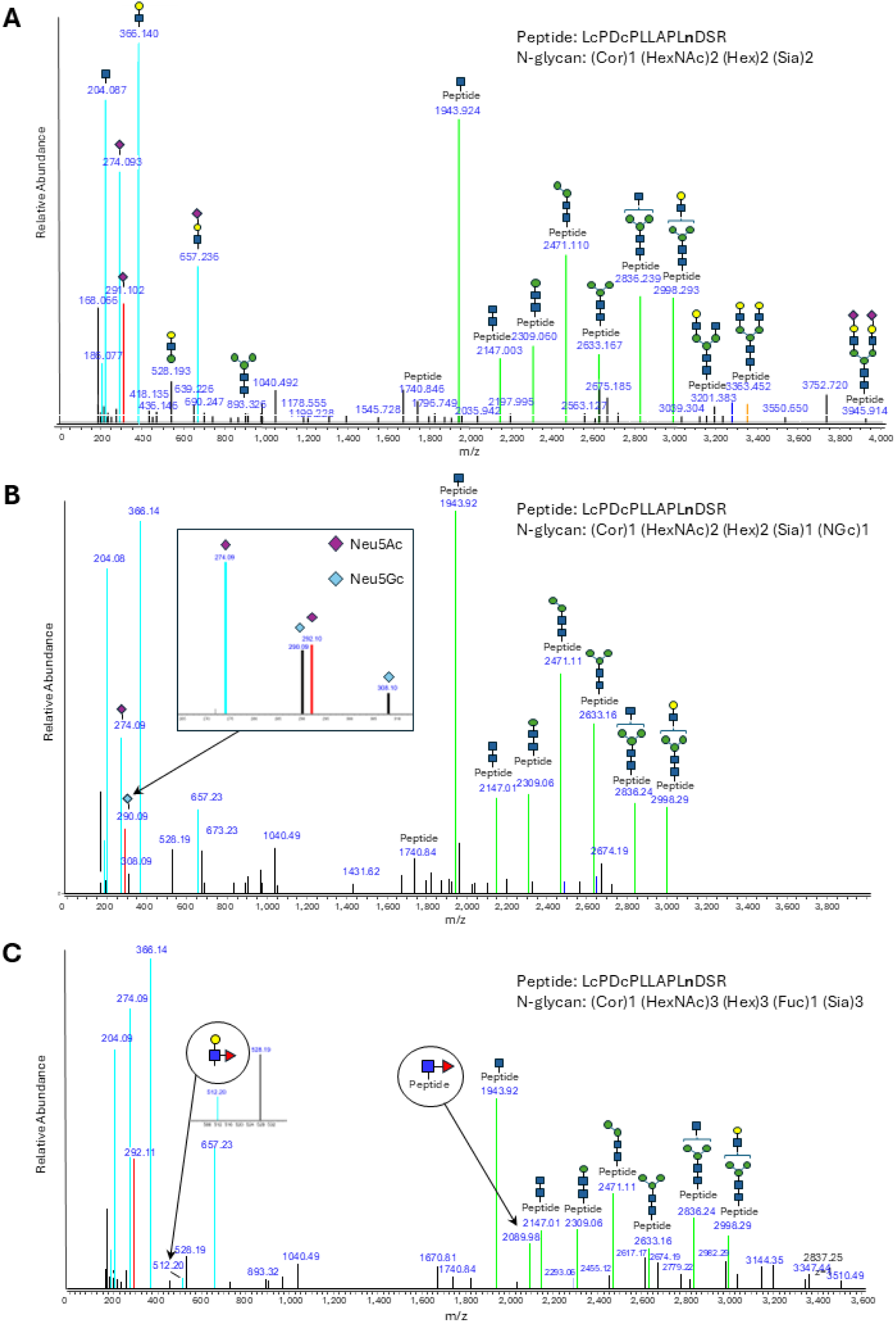
Validation of assigned intact sialylated N-glycopeptide from fetuin using GPMAW glyco-search. (A) The N-glycan composition *(Cor)1 (HexNAc)2 (Hex)2 (Sia)2* was assigned to the fetuin glycosite N156 on the peptide LcPDcPLLAPL<u>n</u>DSR using GPMAW glyco-search. The deconvoluted spectrum, used for the identification, is annotated in colors representing glycan oxonium ions (cyan), Neu5Ac-specific oxonium ions (red), and glycopeptide Y-ions (green). (B) Adding a single Neu5Gc sialic acid in parallel with Neu5Ac during GPMAW glyco-search, the N-glycan composition *(Cor)1 (HexNAc)2 (Hex)2 (Sia)1 (NGc)1* was assigned to the fetuin glycosite N156 on the peptide LcPDcPLLAPL<u>n</u>DSR. Here, Sia denotes the Neu5Ac sialic acid variant (purple diamond), and NGc denotes the non-human Neu5Gc sialic acid variant (blue diamond). The presence of Neu5Gc was supported by diagnostic oxonium ions at m/z 290.09 and 308.1. (C) The N-glycan *(Cor)1 (HexNAc)3 (Hex)3 (Fuc)1 (Sia)3* was assigned to the fetuin glycosite N156 on the peptide LcPDcPLLAPL<u>n</u>DSR. Here, the annotated MS/MS spectrum supports *c*ore fucosylation by the Y1+Fuc fragment, whereas a fucose-containing oxonium ion at m/z 512.2 indicates antenna-associated fucosylated glycan fragments on a co-isolated glycan composition.

This GPMAW glyco-search workflow enables site-specific identification of intact N-glycopeptides together with characterization of glycan compositional heterogeneity at individual glycosites. However, the assigned glycan compositions do not resolve glycosidic linkages, branching patterns, or structural isomers (e.g., α2,3- and α2,6-linked sialic acids).

In the present study, the GPMAW glyco-search workflow was combined with selective TiO_2_ enrichment of sialylated N-glycopeptides. By integrating analyses of intact sialylated N-glycopeptides with their corresponding deglycosylated peptides, the workflow provides site-specific glycan compositional information that is lost when analyses are restricted to deglycosylated peptides alone.

### Workflow optimization using the glycoprotein fetuin

The workflow was optimized using bovine fetuin, a well-characterized glycoprotein commonly used as a standard for method development due to its known glycosites and diverse glycan structures. Fetuin carries both N- and O-linked glycans with a significant portion of them being sialylated. Specifically, fetuin contains three N-glycosites at positions N99, N156, and N76 (**Table 1**). Bovine fetuin was digested with trypsin and selective enrichment for sialylated N-glycopeptides was performed using TiO_2_ chromatography. The GPMAW glyco-search workflow (**Figure 1**) was applied to the enriched sialylated N-glycopeptides from fetuin, and candidate intact N-glycopeptides were identified as described above. Candidate N-glycopeptides with confidence scores over 650 were accepted, whereas those with scores between 450 and 650 were manually validated. Validation was conducted manually in the “Graph / evaluation” interface of the GPMAW glyco-search program by verifying the presence of signature fragment ions Y1 and Y5, and associated glycan fragments in the imported MS/MS spectrum (**Figure 2A**). Additionally, oxonium ions specific for sialylated N-glycans (N-glycan specific: m/z 204.09 and 366.14, Neu5Ac specific: m/z 274.09 and 292.1) were found and verified.

**Table 1:**
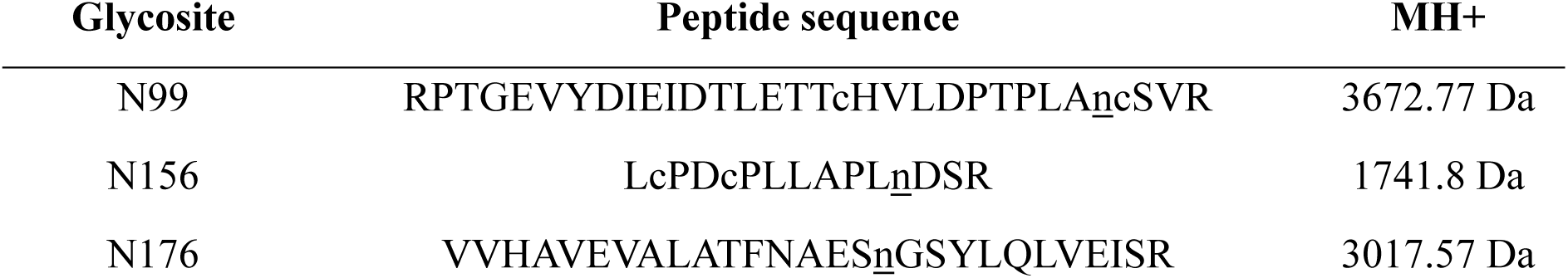
Overview of tryptic N-deglycopeptides identified in bovine fetuin with information on their glycosite, peptide sequence, and MH+. For this study, modifications included carbamidomethylation of cysteines from alkylation with iodoacetamide, and deamidation of asparagine from deglycosylation with PNGase F/A. The modifications are annotated in lower case letters, where deamidation of asparagine is further underlined to highlight the N-glycosite.

Deconvoluting the imported spectra (from the MGF file) in GPMAW increases the number of fragment ions that are assigned during identification and validation with the GPMAW glyco-search program and increases the Y1 ion intensity in case of multi-charged fragment ions from larger N-glycopeptides. Furthermore, the deconvolution results in a better calculation of the score.

**Figure 2A** illustrates the GPMAW glyco-search validation interface using the identification of the N-glycan composition (Core)1(HexNAc)2(Hex)2(Sia)2 assigned to glycosite N156 of the peptide LcPDcPLLAPLnDSR as an example. Here, the deconvoluted MS/MS spectrum is displayed in the GPMAW glyco-search “Graph / evaluation” interface, including color-coded annotation of the peptide Y-ion series (see **Supplementary File S1** for a detailed description).

The glycan composition corresponding to the individual signals are added annually to the figure. Comparison of the theoretical fragment ions with the deconvoluted spectrum confirms the glycopeptide assignment.

The deconvolution of the spectra from the MGF file can be done using various programs and in various ways. The deconvolution in the GPMAW program is described in **Supplementary File S1**. For the specific spectrum shown in **Figure 2A** the efficiency of deconvolution in GPMAW has been compared with the deconvolution performed in the Freestyle v16.1 program (Thermo Fisher Scientific)(**Supplementary Figure S1**). **Figure S1A** shows the GPMAW generated deconvoluted and annotated spectrum, **Figure S1B** shows the same spectrum after deconvolution using Freestyle v16.1, whereas **Figure S1C** shows the corresponding raw MS/MS spectrum without deconvolution. The close agreement between the GPMAW and Freestyle deconvoluted spectra demonstrates that the GPMAW deconvolution performs comparably to Freestyle.

Although most sialylated N-glycopeptides were successfully identified, a small proportion of larger intact glycopeptides could not be validated because of incorrect monoisotopic precursor assignment by the Orbitrap Eclipse. These errors primarily affected low-abundance precursor ions with high m/z values and higher charge states, resulting in mass shifts of +1 or +2 Da. Despite monoisotopic correction during MGF file generation in PD, these errors persisted and prevented correct glycan assignment by high-accuracy mass mapping. This limitation is expected to become increasingly important for complex biological samples containing low-abundance glycopeptides. This limitation was effectively addressed by the subsequent GPMAW glyco-search validation and filtering steps, which removed incorrect glycopeptide assignments caused by erroneous monoisotopic precursor selection.

High-accuracy mass mapping in GPMAW glyco-search performs comprehensive matching against all glycan compositions contained in the selected glycan composition database. Consequently, multiple glycan compositions with similar masses may satisfy the precursor mass tolerance, particularly when monoisotopic precursor assignment is incorrect or mass difference falls within the instrument accuracy. One of the consequences could be that glycan structures containing two fucose residues may be erroneously assigned instead of one Neu5Ac due to an incorrect assignment of the monoisotopic mass. To minimize these misassignments, we limited the high-accuracy mass mapping algorithm to allow a maximum of one fucose residue per intact N-glycopeptide, thereby reducing false-positive assignments to multiply fucosylated glycan compositions. This restriction preludes the identification of glycan compositions containing more than one fucose residue; however, the maximum number of allowed fucose residues can readily be adjusted according to the expected glycan repertoire of the intact N-glycopeptide fragment ion series.

The GPMAW glyco-search workflow in combination with TiO_2_ enrichment identified a total of 21 unique sialylated N-glycan compositions across the three N-glycosites in bovine fetuin were identified, corresponding to 14 on N99, 16 on N156, and 1 on N176. A total of 31 unique intact sialylated N-glycopeptides were identified in fetuin (**Supplementary Data S2**). The total ion chromatogram of the intact sialylated N-glycopeptides analyzed by LC-MS/MS is shown in **Figure 3** with the sialylated N-glycan compositions mapped to each N-glycosite. This demonstrates the ability of the GPMAW glyco-search tool to quickly and easily characterize the compositional heterogeneity of sialylated N-glycans attached to a single N-glycosite.

**Figure 3:**
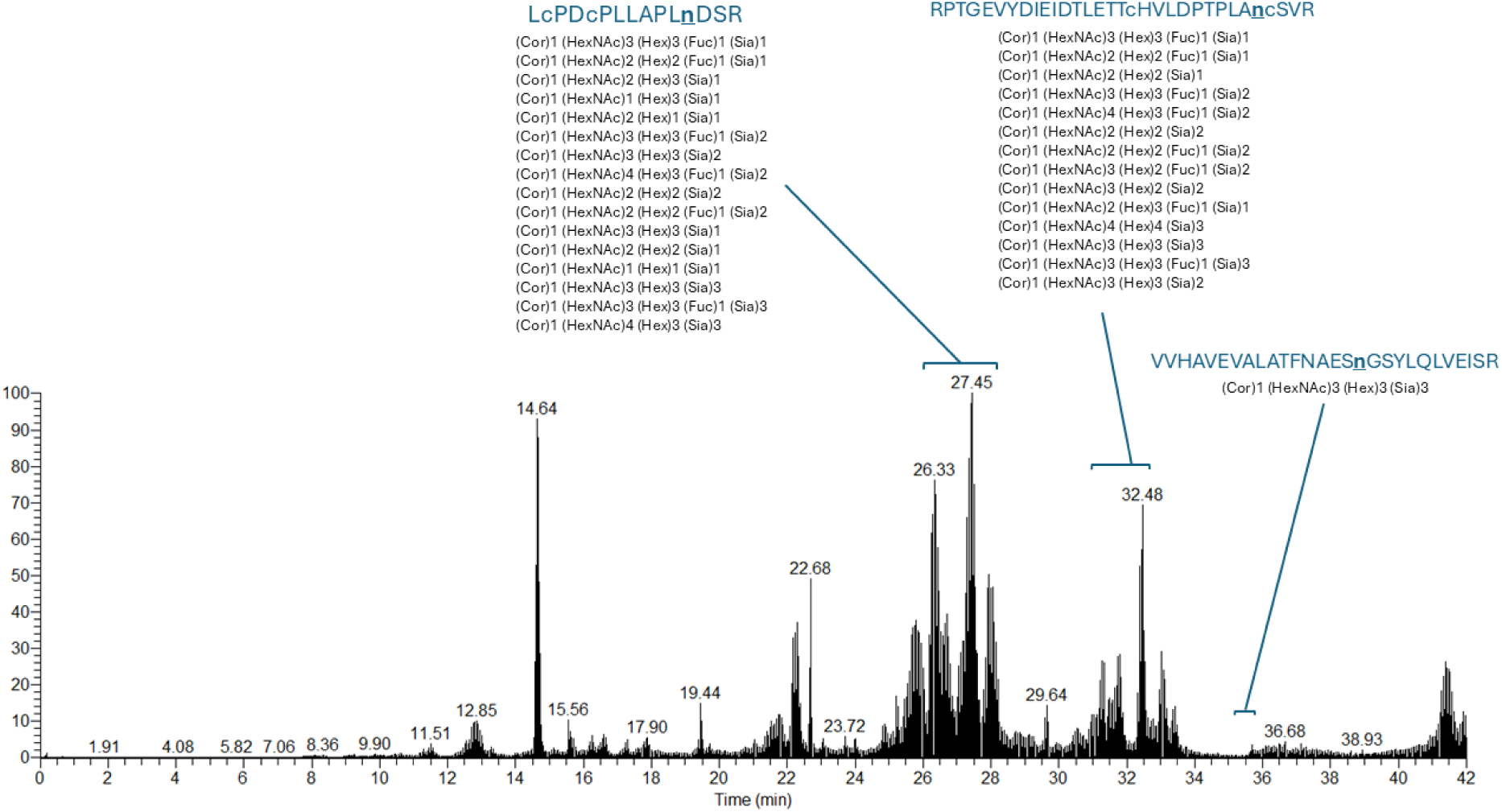
Sialylated N-glycan heterogeneity identified on bovine fetuin. Total ion chromatogram from a 30-minute LC-MS/MS analysis of intact sialylated N-glycopeptides enriched with TiO_2_ chromatography from bovine fetuin. Marked elution regions correspond to intact sialylated N-glycopeptides carrying different N-glycan compositions for the three fetuin N-glycosites, N99, N156, and N176 on peptides RPTGEVYDIEIDTLETTcHVLDPTPLA<u>n</u>cSVR, LcPDcPLLAPL<u>n</u>DSR, and VVHAVEVALATFNAES<u>n</u>GSYLQLVEISR, respectively.

The standard search configuration in GPMAW glyco-search is optimized for the human sialic acid variant, Neu5Ac. The detection of a single Neu5Gc residue along with Neu5Ac during GPMAW glyco-search can be enabled through the setting that adds a single Neu5Gc parallel with Neu5Ac during the search. In bovine protein samples (and most other non-human proteomes) such adjustments expand the dynamic range of identifiable intact sialylated N-glycopeptides. Addition of a single Neu5Gc in parallel with Neu5Ac yielded a total of 30 unique sialylated N-glycan compositions across 49 unique sialylated N-glycopeptides in bovine fetuin (**Supplementary Data S2**). A total of 15 of the sialylated N-glycan compositions contained both sialic acid variants and 3 of the sialylated N-glycan compositions contained only the Neu5Gc variant. The presence of Neu5Gc was verified by oxonium ions m/z 290.09 and 308.1 and by manual interrogation of the intact N-glycopeptide fragment ion series (**Figure 2B**).

In addition, GPMAW glyco-search can identify core-fucosylation by fragment ions corresponding to the mass of Y1+Fuc or outer arm fucosylation by the presence of the oxonium ion at 512.2 Da (**Figure 2C**). These user-configurable options increase the flexibility of the GPMAW glyco-search workflow, making it well suited for high-throughput identification of intact (sialylated) N-glycopeptides across diverse N-glycoproteomics applications.

### Validation of GPMAW glyco-search on standard glycoproteins

The GPMAW glyco-search workflow was applied to four other commercially available glycoproteins; alpha-1-acid glycoprotein 1 (human), serotransferrin (human), fibrinogen alpha/beta chain (human), and thyroglobulin (bovine). These glycoproteins were analyzed using the same workflow as described for fetuin. The resulting deglycopeptide identifications and corresponding MGF files were subsequently imported into the GPMAW glyco-search tool.

After filtering, scoring and validation of individual intact glycopeptides a total of 147 unique N-glycopeptides were identified on four N-glycosites in alpha-1-acid glycoprotein 1 (N33 (25), N56 (23), N72 (69) and N93 (30)). A total of 116 unique N-glycopeptides were identified on serotransferrin (N432 (25), N491 (9) and N630 (82), one N-glycopeptide on the fibrinogen alpha chain (N686) and 27 N-glycopeptides on the fibrinogen beta chain (N394 (27)). On bovine thyroglobulin the search was performed allowing only Neu5Ac species which resulted in the identification of 160 intact N-glycopeptides (N110 (9), N495 (30), N947 (30), N1140 (24), N1776 (52) and N2251 (15)). An overview of the identified N-glycopeptides is shown in **Table 2** and the lists of identified N-glycopeptides from each protein are in **Supplementary Data S3**. The table shows that 451 unique N-glycopeptides were identified from the four proteins and more than 95.8% carried Neu5Ac (432). This, together with the analysis of bovine fetuin above, illustrates the efficiency of identifying (sialylated) N-glycopeptides using the GPMAW glyco-search workflow.

**Table 2:** Overview of unique intact N-glycopeptide identifications across 4 model glycoproteins; alpha-1-acid glycoprotein 1 (human), serotransferrin (human), fibrinogen alpha/beta chain (bovine), and thyroglobulin (human), using GPMAW glyco-search. Search parameter requirements include the presence of the Y1 fragment ion, Y5 fragment ion, and the HexNAc oxonium ion. Further filtration required a Y1 fragment ion peak intensity of at least 20000, a GPMAW score of at least 650, and no oligo-sialylation (number of Hex ≥ number of Sia). Summarization for all 4 glycoproteins include protein accessions (UniProt), glycosites, number of identified intact N-glycopeptides, number of identified intact sialylated N-glycopeptides, and total number of isoforms. The full list of intact N-glycopeptide identifications in the 4 model glycoproteins can be found in Supplementary Data S3.

| Glycoprotein | Protein accession | Glycosite | # intact N-glycopeptides | # intact sialylated N-glycopeptides |
| --- | --- | --- | --- | --- |
| Alpha-1-acid glycoprotein 1 | P02763 | N33 | 25 | 25 |
|  |  | N56 | 23 | 23 |
|  |  | N72 | 69 | 69 |
|  |  | N93 | 30 | 30 |
|  |  | **Total** | **147** | **147** |
| Serotransferrin | P02787 | N432 | 25 | 22 |
|  |  | N491 | 9 | 9 |
|  |  | N630 | 82 | 77 |
|  |  | **Total** | **116** | **108** |
| Fibrinogen alpha chain | P02671 | N686 | 1 | 1 |
| Fibrinogen beta chain | P02675 | N394 | 27 | 24 |
|  |  | **Total** | **28** | **25** |
| Thyroglobulin | P01267 | N110 | 9 | 9 |
|  |  | N495 | 30 | 30 |
|  |  | N947 | 30 | 26 |
|  |  | N1140 | 24 | 24 |
|  |  | N1776 | 52 | 49 |
|  |  | N2251 | 15 | 14 |
|  |  | **Total** | **160** | **152** |

### Large-scale identification of intact sialylated N-glycopeptides in depleted human plasma using the GPMAW glyco-search workflow

To evaluate our high-accuracy mass mapping strategy using the GPMAW glyco-search workflow, a total of 40 µL human plasma was depleted for its 14 highest-abundant proteins, using the top 14 antibody depletion kit from Thermo Scientific. Based on the average protein concentration in human plasma (approximately 70 µg/µL) and that the High-Select Top14 Abundant Protein Depletion Resin removes 95% of the total protein in plasma, approximately 140 µg protein should be left after depletion of 40 µL human plasma. Following tryptic digestion of the depleted plasma, sialylated N-glycopeptides were enriched using TiO_2_ chromatography and subsequently used for fractionation into 12 concatenated fractions using high pH RP fractionation (**Supplementary Figure S2** (LC UV Chromatogram)).

After high pH RP fractionation, 1/3 of the material from each of the 12 fractions were subjected to deglycosylation using PNGase F/A and sialidase A. A list of deglycosylated N-linked glycopeptides was generated for each high pH RP fraction after searching in PD using the SEQUEST HT search engine. The remaining 2/3 of the material of each fraction was analyzed twice on the Orbitrap Eclipse Tribrid MS instrument using 60 min LC gradients. The first LC-MS/MS analysis was performed using HCD NCE of 25% and the second LC-MS/MS analysis was performed using sceHCD with NCE of 25%, 28% and 32%. All raw data files from the intact N-glycopeptide analyses were converted to MGF files in PD and subsequently searched individually in GPMAW glyco-search against the deglycopeptide list corresponding to the given high pH RP fraction. Candidate N-glycopeptides were filtered for a Y1 fragment ion peak intensity above 20k, the presence of Y5 and removal of duplicate hits based on peptide sequence and glycan composition. GPMAW glyco-search further restricted glycan assignments to compositions that are structurally consistent with the number of available antennae. Consequently, glycan compositions requiring α2,8-linked oligo-/polysialic acid structures were not considered (e.g., α2,8 di- and tri-sialylation). Manual inspection of several thousand spectra did not reveal convincing identifications or diagnostic fragment ions supporting α2,8-linked oligo-sialylation in human plasma, suggesting that such structures are either absent or extremely rare in the analyzed samples. We therefore filtered glycan compositions that had more sialic acids than terminal antennae structures. Finally, candidate hits with a score below 450 were removed. For all high pH RP fractions the candidate N-glycopeptides with a score between 450 and 650 were manually inspected and false hits were removed to generate final lists of intact N-glycopeptides. The result of the identification of intact N-glycopeptides from the 12 high pH RP fractions using the GPMAW glyco-search workflow is shown in **Figure 4A** and the lists in **Supplementary Data S4**. On average, we identified 541 intact N-glycopeptides in each high pH fraction of which an average of 506 contained at least one Neu5Ac, corresponding to an average sialic acid enrichment efficiency of 93.6% which correlates with our previous findings [16]. In total, the GPMAW glyco-search workflow, which was performed on 1/3 of the high pH RP purified sample (corresponding to a starting material around 50 µg protein before TiO_2_), identified a total of 2855 unique N-glycopeptides where 2596 carried at least one sialic acid, corresponding to 91% (**Figure 4D**). Analysis of the 12 high-pH fractions using HCD at 25% NCE resulted in slightly fewer identified intact sialylated N-glycopeptides than sceHCD, suggesting that sceHCD is the preferred fragmentation method for this workflow (data not shown).

**Figure 4.**
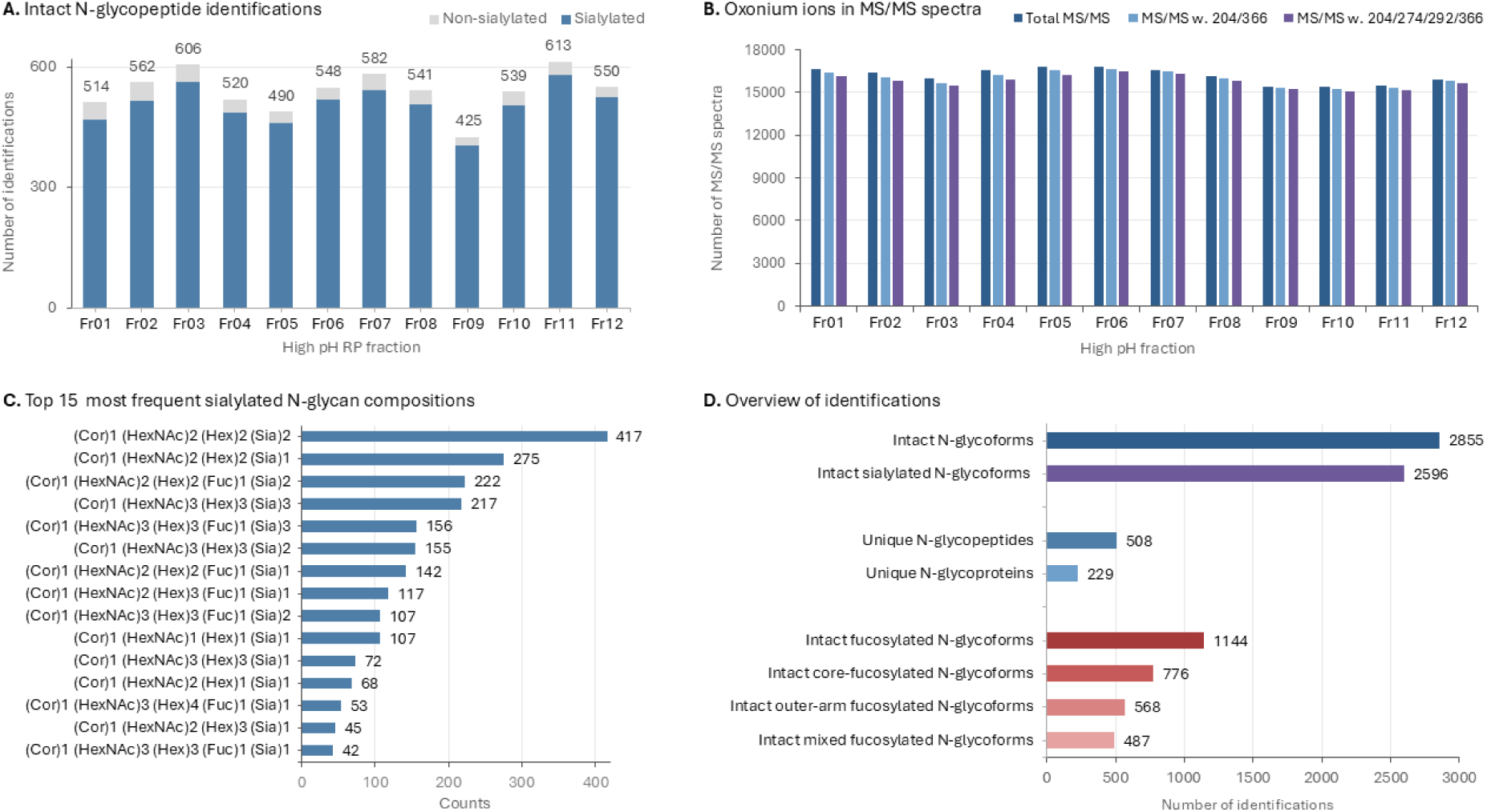
Large-scale identification of intact N-glycopeptides from depleted high-pH reversed-phase fractionated human plasma using GPMAW Glyco-search. (**A**) Number of intact N-glycopeptides identified in each of the 12 high-pH reversed-phase fractions, separated into sialylated and non-sialylated N-glycopeptides. (**B**) Distribution of total MS/MS spectra, spectra containing N-glycan-associated oxonium ions (*m/z* 204.09 and 366.14), and spectra additionally containing Neu5Ac-associated oxonium ions (*m/z* 274.09 and 292.10) identified by filtering for the indicated oxonium ions using the GPMAW MGF File Handling module. (**C**) The 15 most frequently identified sialylated N-glycan compositions across all fractions after removal of redundant glycopeptide identifications. (D) Summary of identified intact N-glycoforms, unique N-glycopeptides, unique N-glycoproteins, and fucosylated N-glycoforms, including core-, outer-arm-, and mixed-fucosylated species.

Using the “MGF File Handling” module in GPMAW, the total number of MS/MS spectra was determined for each of the 12 high pH fractions (MGF files), together with the number of MS/MS spectra containing the four diagnostic glycan oxonium ions at m/z 204.09 (HexNAc), 366.14 (HexNAc+Hex), 274.09 (dehydro-Neu5Ac), and 292.1 (Neu5Ac). Based on these calculations, an average of 98.82% of all MS/MS spectra across the 12 fractions contained the diagnostic glycan oxonium ions at m/z 204.09 and 366.14. Of these glycan-positive spectra, more than 98% also contained the characteristic sialic acid oxonium ions at m/z 274.09 and 292.1. (**Figure 4B**). This illustrates a very high enrichment efficiency for sialylated N-glycopeptides using our optimized TiO_2_ chromatography method. The small discrepancy between the number of identified intact sialylated N-glycopeptides (94%) and the percentage of spectra containing the sialic acid oxonium ions (98%) could be due to co-isolation in the precursor ions selection process where we used 1 Da windows.

Human plasma contains a high protease activity and therefore we performed a search of the deglycopeptides obtained from high pH RP fraction 1 and 2 in PD using enzyme set to “semi-trypsin”. Here we identified almost twice as many unique N-deglycopeptides. Using GPMAW glyco-search with the semi-tryptic peptide lists for the two fractions we identified 30% more intact N-glycopeptides (779 and 804 compared to 553 and 564, respectively (data not shown)) with approximately 93.5% sialylated N-glycopeptides. Extrapolating this to all 12 fractions would correspond to more than 3600 unique intact N-glycopeptides identified from 50 µg depleted plasma if searches were performed with semi-trypsin.

A total of 83 unique N-glycan-compositions were identified in the intact N-glycopeptide list with 61 compositions carrying at least one sialic acid. A summary of the 15 most abundant sialylated N-glycan-compositions identified in depleted plasma (approx. 85% of total sialylated N-glycopeptide identifications) is illustrated in **Figure 4C**. The most prevalent sialylated N-glycan-composition identified in depleted human plasma was (Cor)1 (HexNAc)2 (Hex)2 (Sia)2, occurring on 16.1% of the identified sialylated N-glycopeptides, followed by (Cor)1 (HexNAc)2 (Hex)2 (Sia)1 with 10.6% and (Cor)1 (HexNAc)2 (Hex)2 (Fuc)1 (Sia)2 with 8.6%. Despite using enriched N-glycopeptide originating from only approximately 50 µg of depleted plasma as starting material (1/3 of the original 140 µg enriched material), the GPMAW glyco-search workflow identified 2855 intact N-glycopeptides (**Figure 4D**). A recent large-scale study employing HILIC enrichment combined with spectral library searching identified 526 N-glycoproteins, 1036 N-glycosites, 22677 intact N-glycopeptides, and 738 glycan compositions from human serum, representing one of the deepest serum N-glycoproteomic datasets reported to date [42]. However, the study was designed for comprehensive N-glycoproteome analysis rather than selective characterization of sialylated N-glycopeptides, and the authors do not report how many of the identified N-glycopeptides contained sialic acids. In addition, this analytical depth was achieved using an extensive workflow based on approximately 4 mL pooled human serum (20 × 200 µL; approximately 272 mg total protein), followed by ACN fractionation, repeated HILIC enrichment, PNGase F-assisted spectral library generation, extensive high pH RP fractionation, and multiple LC-MS/MS analyses. In contrast, our workflow specifically enriches sialylated N-glycopeptides directly from substantially smaller sample amounts using a considerably simpler analytical strategy while still achieving deep coverage of the serum sialylated N-glycoproteome. The amount of starting material has a significant influence on the number of N-glycopeptides identified in these studies. Earlier investigations targeting sialylated N-glycopeptides in human plasma reported 22 and 61 identified sialylated N-glycans [43, 44]. A more recent study from 2013 employed TiO₂-enrichment to profile sialylated N-glycopeptides in human plasma, identified a total of 982 sialylated glycosites in 413 glycoproteins [45].

The identification of more than 2800 intact sialylated N-glycopeptides from approximately 50 µg of depleted human plasma demonstrates the high analytical sensitivity and broad applicability of the GPMAW glyco-search workflow for large-scale N-glycoproteomics.

### Identification of sialylated N-glycopeptides in raw human plasma

Next, we aimed to apply our GPMAW glyco-search workflow on a more complex sample, non-depleted human plasma. In three separate replicates we subjected 1 µL of human plasma to the workflow described in **Figure 1**. After enrichment of sialylated N-linked glycopeptides, 90% of the samples were analyzed by LC-MSMS using a two-hour chromatographic gradient using sceHCD fragmentation (25, 28, 32% NCE) and the last 10% of each replicate were subsequently deglycosylated and used for identification of the corresponding N-deglycopeptides. The enrichment efficiency was evaluated for each replicate using the MGF File Handling module in GPMAW on the MGF files from the intact N-glycopeptide LC-MSMS analysis (**Figure 5A**). On average, more than 40000 MS/MS spectra were acquired per analysis. Of these, around 39000 (>95%) contained the diagnostic glycan oxonium ions at m/z 204.09 and 366.14, indicating the presence of glycopeptides. Furthermore, more than 38000 (>98% of the glycopeptide spectra) also contained the characteristic sialic acid oxonium ions at m/z 274.09 and 292.10, demonstrating that the vast majority of enriched glycopeptides were sialylated (**Figure 5A**). These results highlight the exceptional selectivity of the TiO₂ enrichment strategy, even for a highly complex biological sample such as human plasma.

**Figure 5.**
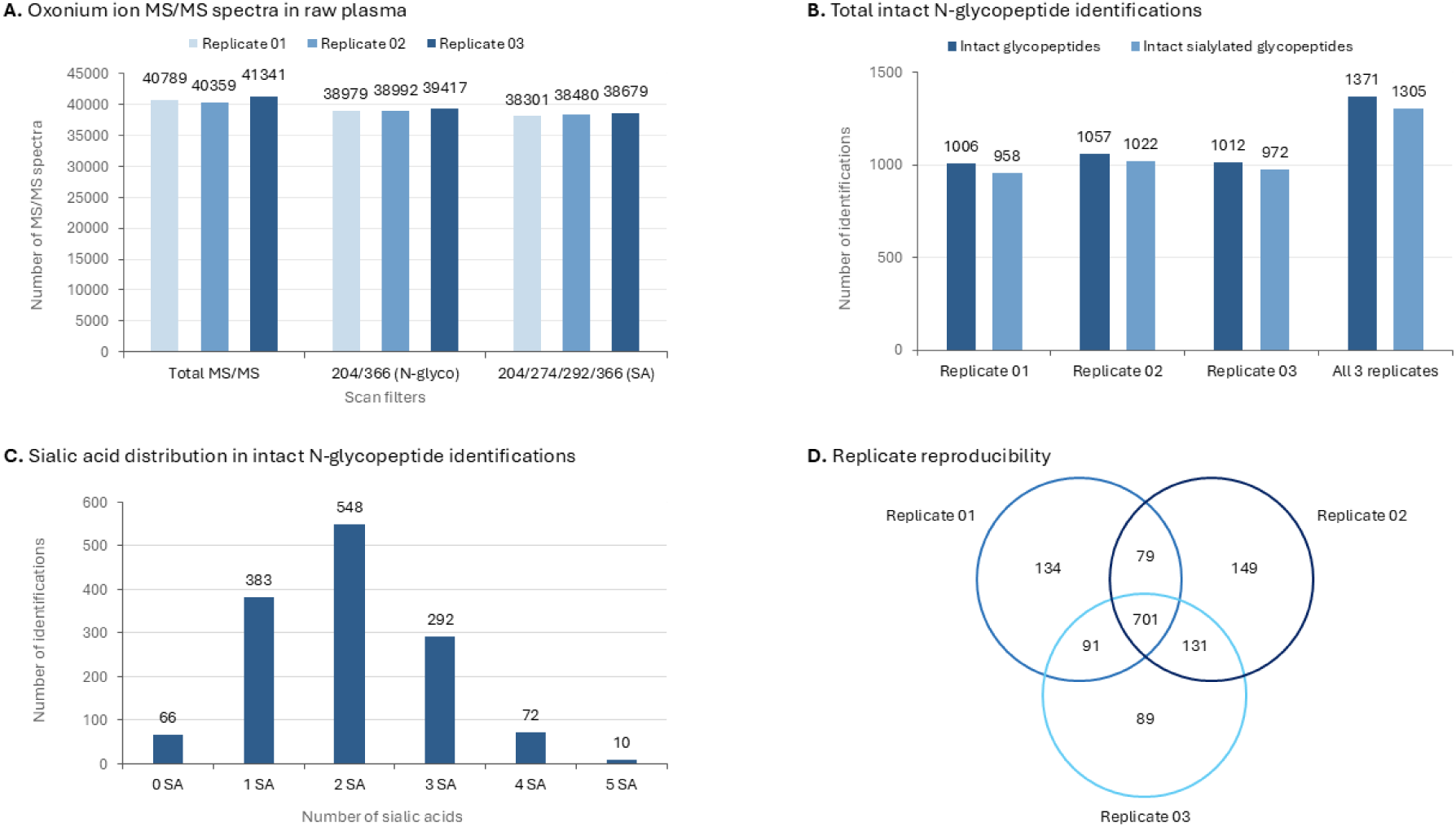
Large-scale identification of intact N-glycopeptides from 1 µL human plasma using GPMAW Glyco-search. (**A**) Total number of MS/MS spectra and spectra containing N-glycan-associated oxonium ions (*m/z* 204.09 and 366.14) and Neu5Ac-associated oxonium ions (*m/z* 274.09 and 292.10) across three replicates following TiO₂ enrichment. (**B**) Number of unique intact N-glycopeptide and intact sialylated N-glycopeptide identifications obtained in each replicate and across all three replicates. (**C**) Distribution of the number of sialic acid residues identified per intact N-glycopeptide. (**D**) Venn diagram illustrates the overlap of intact N-glycopeptide identifications among the three replicates.

The GPMAW glyco-search workflow identified an average of 1025 unique fully tryptic N-glycopeptides across the three biological replicates (1006, 1057, and 1012, respectively) (**Figure 5B**). Of these, an average of 984 were sialylated (958, 1022, and 972, respectively), corresponding to an enrichment specificity of more than 95%. Combining the three replicates yielded a total of 1371 unique N-glycopeptides, of which 1305 were sialylated (**Figure 5B**), where the glycan compositions carrying 2 sialic acids were the most dominant, similar to the results from the depleted plasma (**Figure 5C**). Evaluation of the glycan compositions revealed that 541 were identified with one fucose (core and/or arm) corresponding to 39.4% (**Supplementary Data S6**).

The reproducibility of the GPMAW glyco-search workflow is shown in **Figure 5D**. More than 50% of all identified N-glycopeptides were consistently observed in all three replicates, whereas more than 73% were identified in at least two of the three replicates. Considering that the analyses were performed on an Orbitrap Eclipse using sceHCD fragmentation, relatively long ion injection times, and a comparatively slow MS/MS acquisition rate, this level of overlap demonstrates good analytical reproducibility. At the same time, the incomplete overlap between replicates indicates substantial precursor under-sampling, reflecting the high complexity of the plasma sialylated N-glycoproteome despite the use of a two-hour LC gradient. This is supported by the extensive experimental variability reported in the HUPO Human Glycoproteomics Initiative benchmark study [30], in which substantial variation in identified glycopeptides was observed even when identical LC-MS/MS datasets were analyzed by multiple expert groups using different search strategies.

### Benchmarking GPMAW glyco-search to other N-glycoproteomic search engines

The LC-MS/MS results generated from the triplicate 1µL human plasma was used to evaluate our GPMAW glyco-search tool against other established glycoproteomic search engines: Byonic (v2.10.47, Protein Metrics) [46], MSFragger-Glyco [47], and Glyco-Decipher [37]. The increased sceHCD NCEs for the analyses (25%, 28%, 32%) were originally chosen in order to increase peptide backbone fragmentation and avoid bias against search engines that rely strongly on peptide backbone fragmentation for N-glycopeptide identification.

Byonic performs a peptide-centric database search in which glycans are treated as variable modifications on candidate peptides, and experimental MS/MS spectra are matched against theoretical peptide and glycopeptide fragment ions [46, 48]. MSFragger-Glyco employs a fast open-search strategy to identify candidate peptide backbones and assign glycans by matching the precursor mass offset to entries in a glycan database, supported by diagnostic glycan-derived fragment ions [47, 49]. Glyco-Decipher uses a glycan database-independent, peptide-centric approach in which peptide backbones are inferred from *in silico* deglycosylated spectra and shared peptide fragmentation patterns, followed by assignment of the attached glycan mass [37]. In contrast, GPMAW glyco-search uses high-accuracy mass mapping between intact N-glycopeptide precursor masses and experimentally identified deglycopeptide masses to assign candidate glycan compositions from a user-defined glycan database. Candidate assignments are subsequently validated by the presence of diagnostic glycan fragment ions, including characteristic peptide Y-ions.

To ensure comparable protein search spaces across the search engines, the protein accession numbers from the list of identified deglycopeptides used in the GPMAW glyco-search were used to retrieve the corresponding protein FASTA sequences from UniProt. This FASTA file was subsequently used as the protein database for Byonic, MSFragger-Glyco, and Glyco-Decipher. Precursor monoisotopic assignment can be difficult, as previously mentioned. Each search engine uses different approaches to best correct for this. Byonic allows post-search correction of the assigned precursor mass when it corresponds to an isotope peak instead of a monoisotopic peak. MSFragger-Glyco allows isotope correction during the search by defining a range for allowed precursor monoisotopic assignment errors. Glyco-Decipher itself assigns the precursor by evaluation on isotope and elution pattern during the search. To make a fair comparison we introduced a new feature in GPMAW glyco-search allowing searches with −1, 0 and +1 correction of the precursor ions in the MGF files. Thus, precursor isotope correction was set to allow −1, 0 and +1 for Byonic, MSFragger-Glyco, and GPMAW glyco-search for the most optimal comparison of the search engines.

Each triplicate was searched independently with each search engine, and identifications were subsequently annotated to a common notation based on peptide sequence and N-glycan composition for unique N-glycopeptide assignments. The number of unique N-glycopeptide identifications obtained in each individual replicate was compared for GPMAW, Byonic, MSFragger-Glyco, and Glyco-Decipher (**Figure 6A**). For GPMAW glyco-search the inclusion of −1, 0 and +1 correction of the precursor isotope resulted in more than 16% more unique N-glycopeptides compared with using only 0. GPMAW glyco-search identified 1183, 1215, and 1194 intact N-glycopeptides across the triplicates, yielding 1615 unique N-glycopeptides of which 1523 (94.3%) were sialylated (**Supplementary Data S7**). Byonic identified 1079, 1058, and 1055 intact N-glycopeptides across the triplicates (PEP 2D < 0.001, no manual validation), yielding a total of 1543 unique intact N-glycopeptides of which 1428 (92.6%) contained at least one Neu5Ac (**Supplementary Data S8**). MSFragger-Glyco identified 1111, 1069, and 1069 intact N-glycopeptides across the triplicates (glycan q-value < 0.001, no manual validation), yielding 1540 unique N-glycopeptides of which 1427 (92.7%) were sialylated (**Supplementary Data S9**). Glyco-Decipher identified 886, 1013, and 1030 intact N-glycopeptides across the triplicates (glycan FDR < 0.001, no manual validation), yielding 1561 unique N-glycopeptides of which 1485 (95.1%) were sialylated (**Supplementary Data S10**). The high proportion of intact sialylated N-glycopeptide identifications by all 4 search engines is consistent with the enrichment efficiency by TiO_2_ chromatography previously mentioned, indicating the differences in identifications between the search engines were not caused by differences in overall sialylated N-glycopeptide identification. All four glycoproteomic search engines identified over 1500 unique N-glycopeptides across the three replicates, indicating that the identification coverage of GPMAW glyco-search is similar to already established N-glycoproteomic tools.

**Figure 6.**
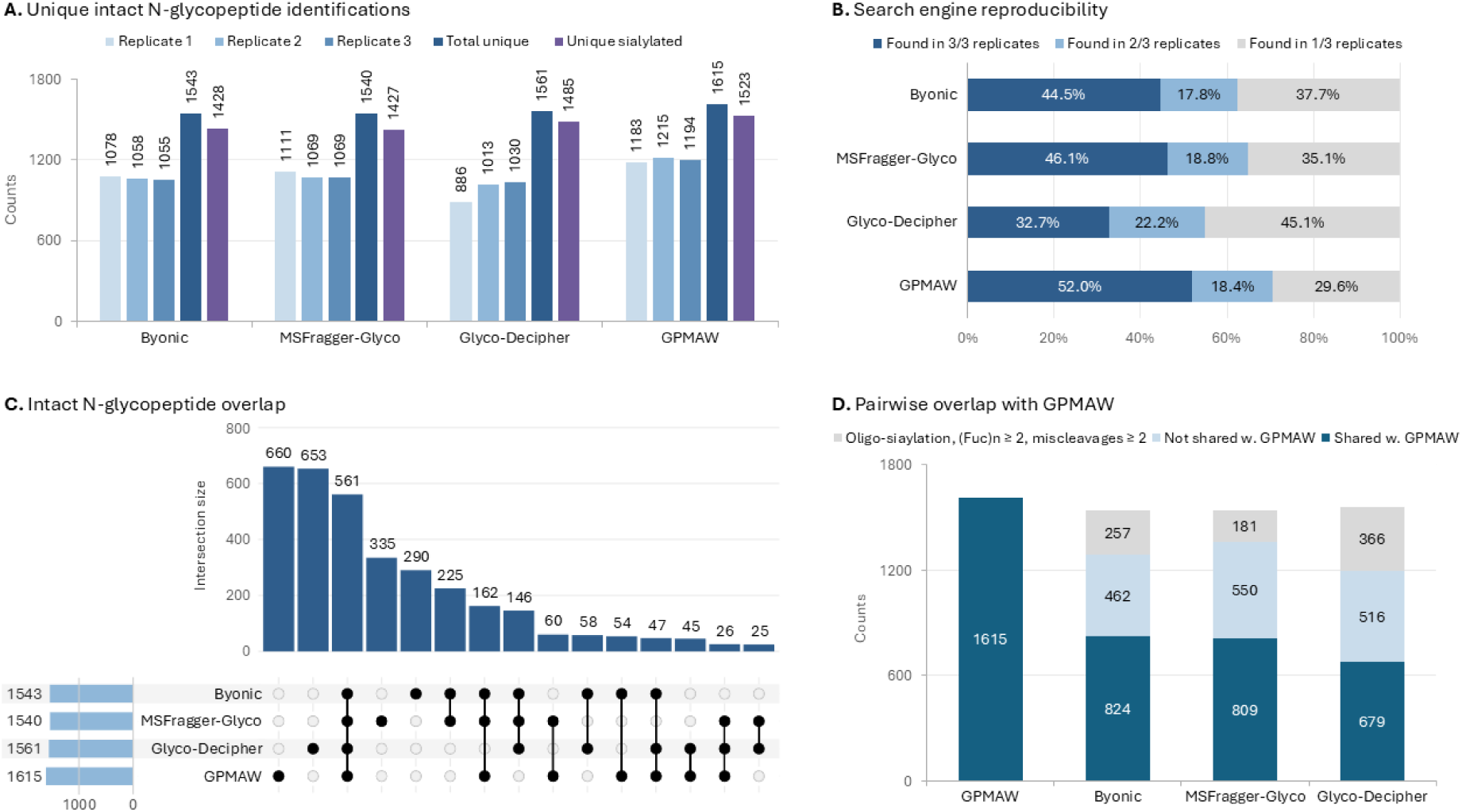
Comparison of intact N-glycopeptide identifications across glycoproteomic search engines. Triplicate datasets of TiO₂-enriched intact sialylated N-glycopeptides from 1 µL human plasma were analyzed using Byonic, MSFragger-Glyco, Glyco-Decipher and GPMAW Glyco-search. (**A**) Number of unique intact N-glycopeptide identifications in each replicate, together with the total number of unique intact N-glycopeptides and unique sialylated N-glycopeptides identified by each search engine. (**B**) Reproducibility of each search engine, shown as the percentage of intact N-glycopeptides identified in three, two or one of the three replicate analyses. (**C**) UpSet plot showing the overlap of non-redundant intact N-glycopeptide identifications among the four search engines. (**D**) Pairwise comparison of GPMAW Glyco-search with each of the other search engines. Grey bars represent identifications outside the GPMAW search space (glycopeptides containing oligo-sialylation, ≥2 fucose residues or ≥2 missed cleavages), light blue bars indicate identifications unique to the respective search engine, and dark blue bars indicate identifications shared with GPMAW Glyco-search.

Next, we assessed the reproducibility within each search engine. Overlap of unique identifications from each of the triplicates, based on peptide sequence and N-glycan composition, was individually assessed for each search-engine and sorted whether they were found in all 3 replicates, 2 of the 3 replicates, or in only 1 of the replicates (**Figure 6B**). This allowed us to evaluate whether each search engine produced consistent N-glycopeptide identifications across repeated analyses of the same sample. Although all four search engines had a similar number of total unique N-glycopeptide identifications, the reproducibility of identifications between the triplicates for each search engine varied considerably. For Glyco-Decipher a large portion of the unique N-glycopeptides were found in only 1 of the 3 replicates (45.1%). In comparison, 37.7%, 35.1%, and 29.6% were found in only one replicate in Byonic, MSFragger-Glyco, and GPMAW glyco-search, respectively. For GPMAW glyco-search only 29.6% of identified N-glycopeptides were observed in a single replicate, whereas 70.4% were identified in at least two of the three replicates. Notably, 52% of all identified N-glycopeptides were consistently detected in all three replicates, highlighting the high reproducibility of GPMAW glyco-search across the replicate analyses (**Figure 6B**).

The overlap between N-glycopeptide identifications obtained with the four search engines was evaluated to determine the extent to which they identified the same N-glycopeptide population (**Figure 6C**). A total of 561 intact N-glycopeptides were identified by all four search engines, representing a robust core dataset that was consistently detected irrespective of the search strategy. However, each search engine also reported unique or partially overlapping N-glycopeptides, that could reflect the fundamentally different search strategies, algorithms and filtering criteria employed for N-glycopeptide identification (**Supplementary Figure S3C**). Using all 4 search engines resulted in the identification of 3347 unique N-glycopeptides from the 3 replicates originating from 1µL plasma.

The large overlap between GPMAW and the three established search engines indicates that most GPMAW identifications are supported by independent algorithms, whereas the additional GPMAW identifications primarily extend rather than replace the common N-glycopeptide repertoire.

However, a detailed manual comparison of the search results showed that a substantial proportion of the N-glycopeptides uniquely reported by Byonic, MSFragger-Glyco and Glyco-Decipher originated from differences in the defined search space rather than fundamentally different interpretation of identical spectra (**Figure 6D and Supplementary Figure S3**). Specifically, 257, 181 and 366 unique N-glycopeptides reported by Byonic, MSFragger-Glyco and Glyco-Decipher, respectively, contained features that were intentionally excluded from the GPMAW glyco-search workflow. These included glycans carrying more than one fucose residue, peptide sequences with two or more missed tryptic cleavages, and glycan compositions containing more sialic acid residues than permitted by the number of glycan antennae, which would require the presence of α2,8-linked oligo-/polysialic acid structures in human plasma (**Supplementary Figure S3A and Figure S3B**).

In the GPMAW glyco-search tool the search was restricted to a maximum of one fucose residue, both because multiply fucosylated N-glycans are generally uncommon in normal human plasma (1.6% dual fucosylated N-glycopeptides) [50] and because glycan compositions containing two fucose residues are nearly isobaric with compositions containing one additional sialic acid. Incorrect monoisotopic precursor assignment may therefore lead to ambiguous glycan composition assignments. Consistent with this, the recent HUPO Human Glycoproteomics Initiative benchmark study identified multiply fucosylated glycopeptides as one of the glycan classes most frequently misassigned by current glycoproteomics search engines [30]. Several of the alternative search engines reported N-glycan compositions containing two to five fucose residues. Given the low reported abundance of multiply fucosylated N-glycans in normal human plasma, together with the ambiguity introduced by near-isobaric glycan compositions and incorrect monoisotopic precursor assignment, these identifications should be interpreted with caution and ideally be supported by manual spectrum validation.

Similarly, glycan compositions requiring α2,8-linked oligo-sialic acid structures were excluded from the GPMAW glyco-search search space. Manual inspection of several thousand annotated MS/MS spectra did not reveal convincing diagnostic fragment ions supporting such structures in human plasma. Furthermore, interrogation of more than 40000 MS/MS spectra per replicate using the GPMAW MGF File Handling module identified only 15 spectra containing the complete set of diagnostic fragment ions expected for these α2,8-linked oligo-sialylated N-glycan structures. Manual inspection of these spectra revealed that only three originated from N-linked glycopeptides, and in all three cases the apparent oligosialylation-specific oxonium ions resulted from low-intensity co-isolated precursor ions rather than the N-glycopeptide itself (data not shown).

Furthermore, GPMAW glyco-search uses experimentally identified deglycopeptides as the basis for glycan assignment rather than searching theoretical protein FASTA databases. The deglycopeptide list is generated by enzymatic deglycosylation followed by database searching and extensive validation. To further increase assignment specificity, the GPMAW glyco-search workflow is restricted to peptides containing a maximum of one missed cleavage, thereby reducing search-space complexity, and minimizing redundant peptide assignments to the same glycosite. In contrast, most conventional N-glycoproteomics search engines perform database searches against the whole protein sequence and typically allow two or more missed cleavages. In addition, we limited our deglycopeptide analysis to select and fragment peptides with minimum 2 charges and above 300 m/z, which would exclude all potential deglycopeptides with a mass below 600 Da. Consequently, a proportion of the differences observed between GPMAW glyco-search and the other search engines reflects differences in the underlying peptide search space rather than glycan assignment alone.

Collectively, these observations suggest that a substantial proportion of the apparent disagreement between search engines reflects differences in search-space definition rather than fundamentally different interpretation of identical glycopeptide spectra.

Overall, GPMAW glyco-search outperformed the three search engines evaluated in this study with respect to confident identification of intact sialylated N-glycopeptides and number of N-glycopeptides identified. During software development, manual evaluation of several thousand annotated spectra using the integrated spectrum viewer demonstrated that identifications with GPMAW scores above 650 are highly reliable and associated with a very low false-positive rate. Unlike conventional search engines, GPMAW glyco-search does not rely on extensive peptide backbone fragmentation but instead used high accuracy MS and MS/MS data and exploits the highly informative glycan Y-ion series together with experimentally defined peptide constraints, making the approach particularly well suited for highly sialylated N-glycopeptides, where peptide backbone fragmentation is frequently limited [16]. In addition, the software provides considerable flexibility through user-defined glycan databases and can readily be extended to include modified glycan compositions, and it contains structural features such as core- and outer-arm fucosylation. Although demonstrated here using sialylated plasma N-glycopeptides, GPMAW glyco-search is not restricted to sialylated species. These characteristics make it a robust and adaptable platform for comprehensive analysis of intact N-glycopeptides across diverse biological samples and glycan classes.

## Conclusion

In this study, we present a robust workflow for the identification and validation of intact sialylated N-glycopeptides that combines highly selective TiO₂ enrichment, dual LC-MS/MS analysis of intact and deglycosylated glycopeptides, and the GPMAW glyco-search platform based on high-accuracy mass mapping. By using experimentally identified deglycopeptides together with experimentally defined search criteria and diagnostic fragment ion validation, the workflow reduces assignment ambiguity while maintaining extensive coverage of the sialylated N-glycoproteome.

Application of the workflow to standard glycoproteins and human plasma demonstrated high enrichment specificity, deep glycoproteome coverage, and excellent reproducibility. Compared with established N-glycoproteomics search engines, GPMAW glyco-search consistently identified more sialylated N-glycopeptides while providing an intuitive platform for interactive spectrum validation and filtering.

GPMAW glyco-search provides a robust and flexible strategy for confident intact N-glycopeptide identification by combining experimentally identified deglycopeptides with stringent glycan assignment and spectrum validation. The modular framework allows straightforward adaptation to different glycomes and future extensions, making the software broadly applicable for N-glycoproteomics beyond the plasma N-sialiome investigated here.

## Data Availability

The MS proteomics data generated in this study, including the raw data, MGF files, and search files have been deposited to the ProteomeXchange Consortium via the PRIDE [35] partner repository with the dataset identifier PXD081982. The dataset is currently accessible to reviewers through the private PRIDE reviewer account and will be made publicly available upon publication.

## Acknowledgements

This research was financially supported by the Independent Research Fund Denmark (Natural Science) (grant no. 2032-00279B), the Villum Centre for Bioanalytical Sciences at SDU and the Danish Agency of Higher Education and Science to establish the PLATO research infrastructure: Danish National Mass Spectrometry Platform for Proteomics and Biomolecular Imaging (grant no. 5229-00012B, www.sdu.dk/PLATO).

This work was funded by the São Paulo Research Foundation – FAPESP (grant nos. 2018/15549-1, 2022/11334-6, and 2020/04923-0 to G.P., 2017/04032-5 and 2021/14751-4 to S.N.M.

## Author contributions: CRediT

**Maria K. Petersen:** Formal analysis, Investigation, Data curation, Writing – original draft, Writing – review and editing, Visualization. **Simon N. Mule:** Formal analysis, Data curation, Writing – review and editing. **Sara E. Lendal:** Investigation, Writing – original draft. **Arkadiusz Nawrocki:** Methodology, Investigation. **Giuseppe Palmisano:** Methodology, Resources, Writing – review and editing. **Peter Højrup:** Methodology, Software, Resources, Writing – review and editing. **Martin R. Larsen:** Conceptualization, Methodology, Resources, Writing – review and editing, Supervision, Project administration, Funding acquisition.

## Supplementary Figures

**Supplementary Figure S1:**
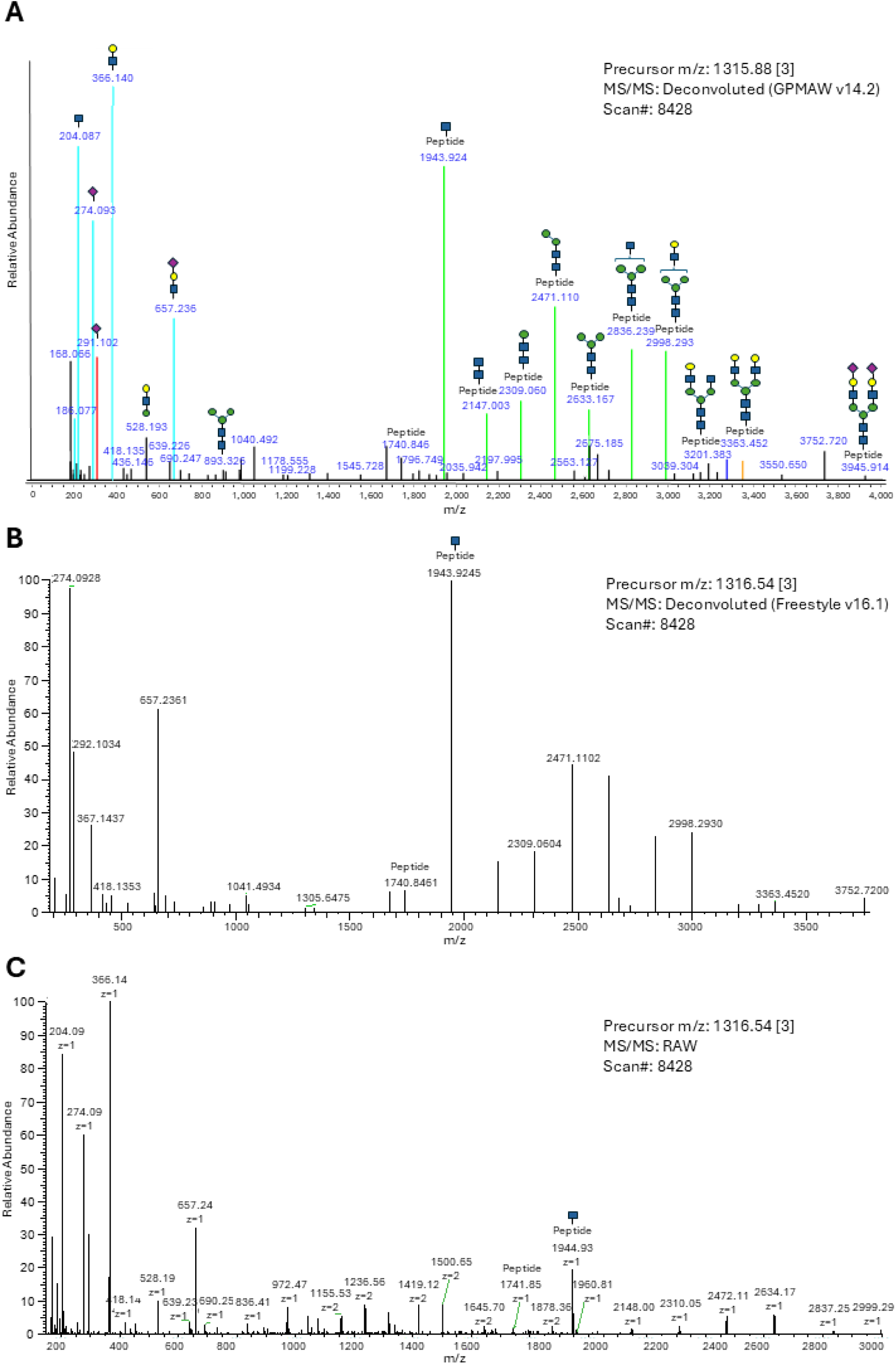
Deconvolution of an intact sialylated N-glycopeptide from fetuin with GPMAW glycosearch. (A) The N-glycan composition *(Cor)1 (HexNAc)2 (Hex)2 (Sia)1* was assigned to the fetuin glycosite N156 on the peptide LcPDcPLLAPL<u>n</u>DSR using GPMAW glycosearch. The MS/MS spectrum from scan #8428 is deconvoluted and annotated in color-labels by GPMAW glyco-search, where glycan oxonium ions are shown in cyan, Neu5Ac-specific oxonium ions in red, and peptide+glycan Y-ions in green. (B) The deconvoluted MS/MS spectrum from the same scan in Freestyle v16.1. (C) Raw MS/MS spectrum of scan # 8428 viewed in Thermo Xcalibur Qual Browser.

**Supplementary Figure S2:**
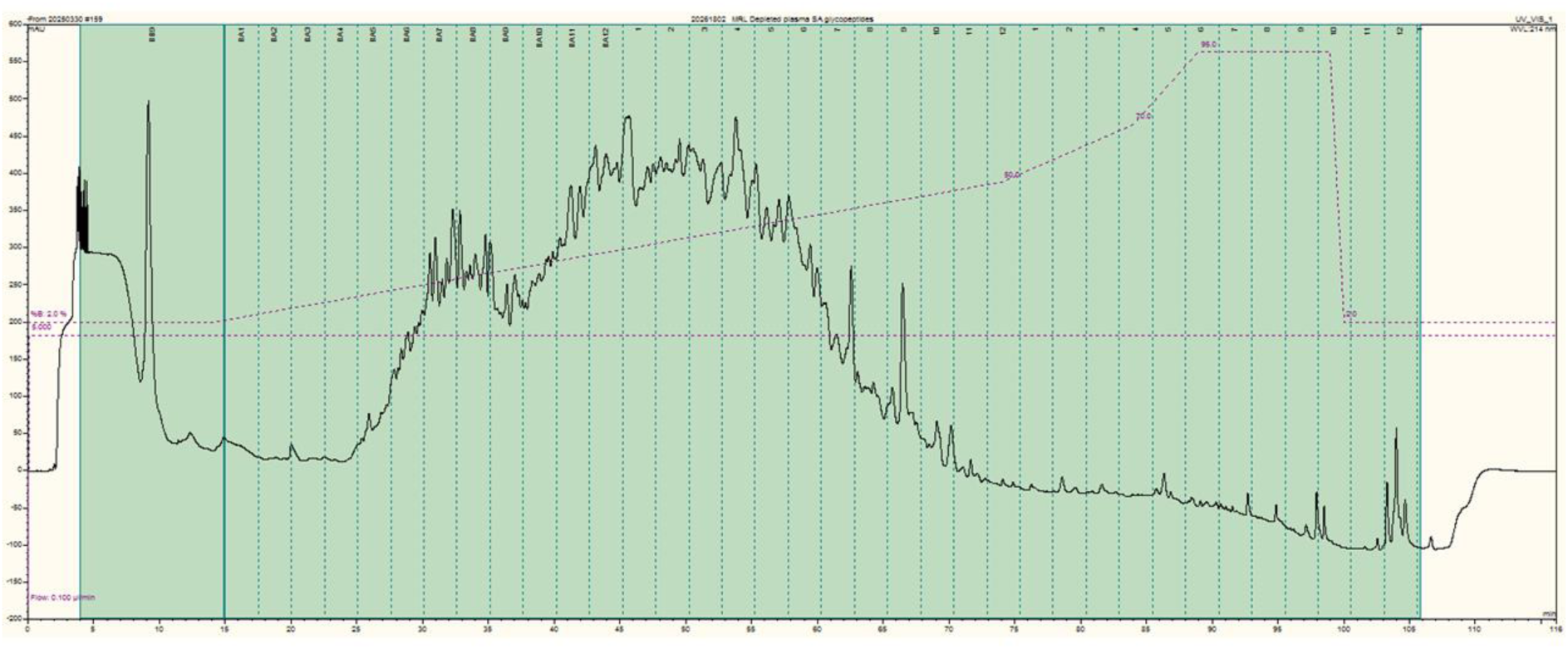
Depleted plasma HpH RP chromatogram. High pH RP LC UV chromatogram of TiO_2_ enriched glycopeptides from depleted plasma. Fractions were obtained by concatenation into 12 unique fractions.

**Supplementary Figure S3.**
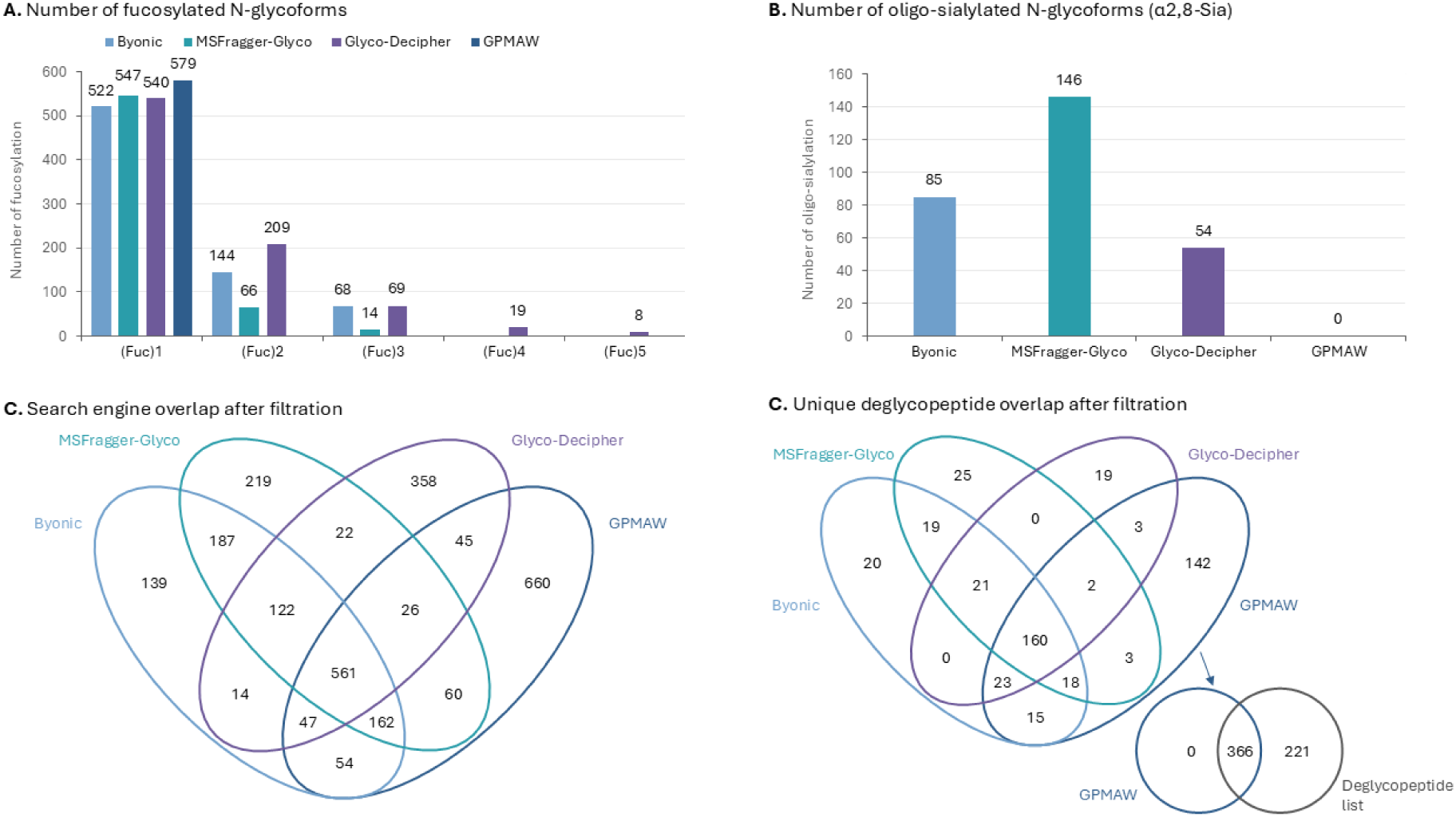
Detailed comparison of glycoproteomic search engine identifications. (**A**) Distribution of identified intact N-glycoforms according to the number of fucose residues assigned per glycan composition. (**B**) Number of identified potential oligo-sialylated N-glycoforms by the programs (glycan compositions containing more sialic acid residues than hexose residues, excluding the N-glycan core). (**C**) Four-way Venn diagram showing the overlap of intact N-glycopeptide identifications among the four search engines following post-search filtration. (**D**) Four-way Venn diagram showing the overlap of peptides from the identified intact N-glycopeptides among the four search engines.

